# “A recombinant protein SARS-CoV-2 candidate vaccine elicits high-titer neutralizing antibodies in macaques”

**DOI:** 10.1101/2020.12.20.422693

**Authors:** Gary Baisa, David Rancour, Keith Mansfield, Monika Burns, Lori Martin, Daise Cunha, Jessica Fischer, Frauke Muecksch, Theodora Hatziioannou, Paul D. Bieniasz, Fritz Schomburg, Kimberly Luke

## Abstract

Vaccines that generate robust and long-lived protective immunity against SARS-CoV-2 infection are urgently required. We assessed the potential of vaccine candidates based on the SARS-CoV-2 spike in cynomolgus macaques (*M. fascicularis*) by examining their ability to generate spike binding antibodies with neutralizing activity. Antigens were derived from two distinct regions of the spike S1 subunit, either the N-terminal domain (NTD) or an extended C-terminal domain containing the receptor-binding domain (RBD) and were fused to the human IgG1 Fc domain. Three groups of 2 animals each were immunized with either antigen, alone or in combination. The development of antibody responses was evaluated through 20 weeks post-immunization. A robust IgG response to the spike protein was detected as early as 2 weeks after immunization with either protein and maintained for over 20 weeks. Sera from animals immunized with antigens derived from the RBD were able to prevent binding of soluble spike proteins to the ACE2 receptor, shown by *in vitro* binding assays, while sera from animals immunized with the NTD alone lacked this activity. Crucially, sera from animals immunized with the RBD but not the NTD had potent neutralizing activity against SARS-CoV-2 pseudotyped virus, with titers in excess of 10,000, greatly exceeding that typically found in convalescent humans. Neutralizing activity persisted for more than 20 weeks. These data support the utility of spike subunit-based antigens as a vaccine for use in humans.

## Background

In less than one year since the emergence of SARS-CoV-2, over 71 million people have been infected worldwide resulting in over 1.3 million deaths^1^. The SARS-CoV-2 pandemic highlights the need for vaccines to slow viral spread, prevent disease and potentially protect against future pandemics. Several SARS-CoV-2 vaccine candidates are currently in Phase III clinical trials or have been approved, including RNA-based and viral vector-based vaccines^2; 3^. While these approaches have yielded promising results, the low temperature storage requirements for certain vaccine candidates create challenges in regards to mass vaccination strategies, particularly in resource limited settings. Recombinant protein-based vaccines or subunit vaccines have been widely used and can be formulated with adjuvants to enhance immunogenicity^4; 5; 6; 7; 8^. Moreover, subunit vaccines can be easily scaled up for mass production while maintaining efficacy and safety.

The goal of an effective vaccine is to produce a long-term protective antibody and or T-cell response. Most potential SARS-CoV-2 vaccines have focused on the viral spike (S) protein. The SARS-CoV-2 spike protein forms a trimer that binds the host cell receptor, angiotensin converting enzyme-2 (ACE2)^9; 10^. The SARS-CoV-2 spike is initially synthesized as a precursor that is ultimately cleaved by cellular proteases to generate the S1 and S2 domains^11^. The S1 domain contains the receptor binding domain (RBD)^12^, that binds ACE2^9; 11; 13; 14^ (Fig. 1A) while the S2 domain contains the stalk portion of the protein and the fusion machinery that enables viral entry, and exhibits higher levels of sequence conservation among coronaviruses (Fig 1A, B)^9; 11; 12^. All SARS-CoV-2 neutralizing antibodies identified thus far in recovered COVID-19 patients bind to the S1 subunit, specifically the N-terminal domain (NTD) or RBD^15; 16; 17; 18; 19,20^ Indeed, many, but not all neutralizing antibodies block the interaction of the RBD with ACE2 receptor. Based on these findings, we cloned and expressed domains of the S1 subunit and tested them as vaccine candidates using a nonhuman primate model. Specifically, we generated distinct antigens that either contained the NTD or the RBD fused to an immunoglobulin Fc domain (NTD-Fc and RBD-Fc).

**Figure 1.**
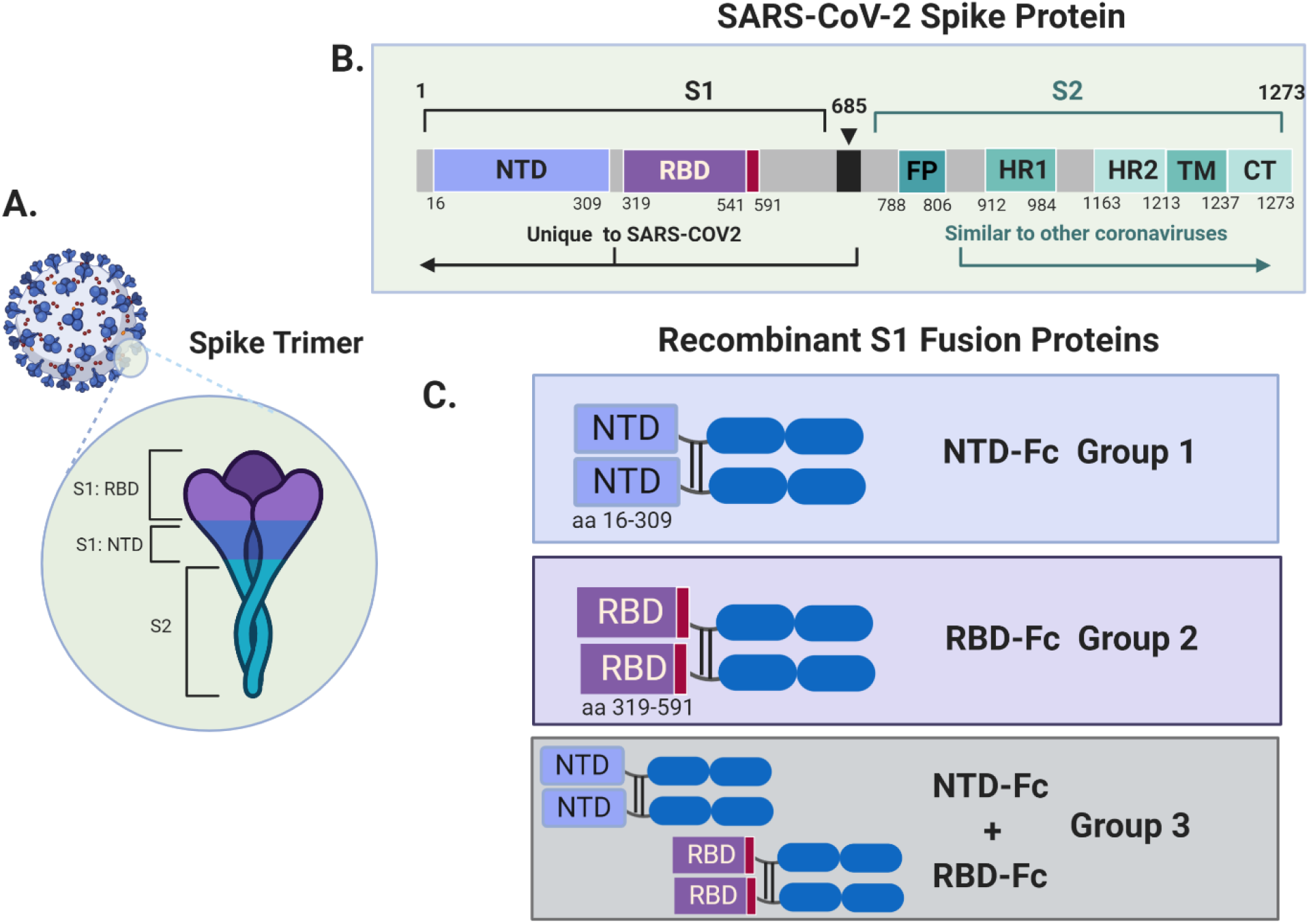
Structure and terminology of SARS-CoV-2 spike constructs. **A**. Major structural features of the SARS-CoV-2 virus include the surface exposed Spike (S) protein, which functions as a trimeric binding partner for the host ACE2 protein. **B**. Structural features of the spike protein. S1: N-terminal half of the protein, cleaved at the S1/S2 site (685) by cellular furin. NTD: N-terminal portion of S1 domain (aa 16-309). RBD: C-terminal portion of the S1 domain includes the receptor binding domain (RBD) that interacts with the human ACE2 receptor (aa 319-541). S2: stalk portion of the spike protein (aa 685-1273), a region with sequence similarity to other related coronaviruses. Contains the fusion peptide (FP), heptapeptide repeat sequences (HR1 and HR2), transmembrane region (TM) and a cytoplasmic domain (CT) ^9; 14^. **C**. Treatment groups with NTD-Fc and RBD-Fc as immunogens. NTD-Fc contains amino acids 16-309 expressed as a fusion protein with human IgG1 Fc fragment. RBD-Fc contains amino acids 319-591 expressed as a fusion protein with human IgG1 Fc fragment. Made with Biorender.com

The nonhuman-primate (NHP) research model has proven invaluable in studies of disease transmission, progression and pathology, and development of therapeutics, vaccines, and diagnostics against SARS-CoV-2^21; 22; 23^. To compare the efficacy of different SARS-CoV-2 spike antigens for use in a potential recombinant subunit vaccine, cynomolgus macaques were immunized with NTD-Fc and or RBD-Fc fusion proteins. To evaluate the antibody response over the course of immunization, a multiplex immunoassay was developed to detect SARS-CoV-2 specific antibodies in nonhuman primates. High spike-binding antibody titers were detected within 2 weeks and through 20 weeks post-immunization, with animals receiving the RBD-Fc generating higher titers. Moreover, immunization with antigens that included the RBD-Fc alone or in combination with the NTD-Fc, resulted in serum with potent viral neutralizing activity. This work demonstrates that vaccination with RBD-containing antigens can elicit a strong and durable neutralizing antibody response in macaques and warrants further investigation for potential use in humans.

## Results

### Production of SARS-CoV-2 NTD-Fc and RBD-Fc fusion proteins

Based on prior studies demonstrating the ability of the spike protein of SARS-CoV and SARS-CoV-2 to elicit a robust immune response^3; 25; 26; 27; 28; 29; 30; 31^, recombinant spike-based proteins were developed for immunization of animal models to evaluate these subunit-based antigens for potential use as vaccine candidates. Structural studies of the spike trimer have demonstrated that it binds the host cell receptor ACE2^9; 10; 17; 32^ (Fig. 1A) and have identified the NTD and RBD as conformationally discrete domains within S1, both of which can be targets of neutralizing antibodies^15; 16; 18^. The N-terminal domain (NTD, aa 16-309) or a region including the RBD (aa319-541) were fused to the C-terminal human IgG1 Fc to generate NTD-Fc and RBD-Fc, respectively (Fig. 1B, C). The RBD-Fc protein includes an additional 50 residues of the C-terminal domain of S1.

### Immunization of macaques

All macaques in this study had a pre-immunization blood draw tested for cross-reactive antibodies to SARS-CoV-2 proteins that could conceivably be elicited by prior exposure to related coronaviruses. Serum was screened using the standard protocol for antibody testing using the CSA: SARS-CoV-2 test, which is a high sensitivity test for identification of antibodies to SARS-CoV-2 spike S1, and S2 subunits, and nucleocapsid antigens. These recombinant antigens are individually immobilized to a specific location or “spot” within a well in quadruplicate, resulting in a multiplex assay for simultaneous detection of antigen-specific antibodies within each well. Signal increases with increasing amount of bound antibody. Results are calculated from the median of four spots in each well, and the mean of 3 replicate wells are graphed. As shown in Figure 2A, all animals had negligible antibody reactivity to SARS-CoV-2spike S1. Overall, all animals were SARS-CoV-2 antibody negative prior to immunization.

**Figure 2.**
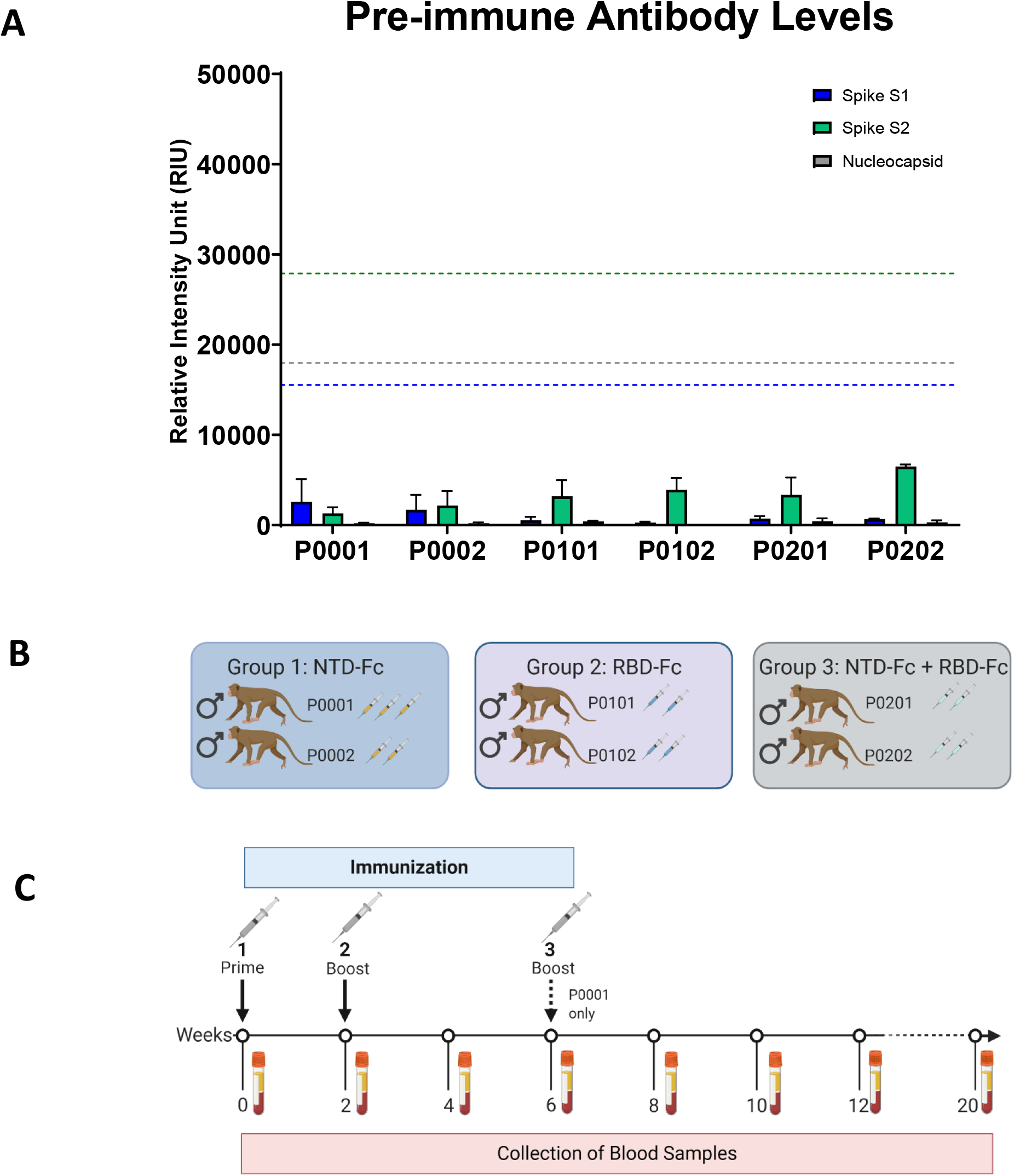
Immunization Groups and Schedule. **A**. Pre-immunization serum screening against SARS-CoV-2 spike S1, S2, and nucleocapsid. Serum from pre-immunization diluted 1:100 (0 weeks) was screened for antibodies using the CSA: SARS-CoV-2 assay for anti-spike S1 (blue), anti-spike S2 (green), and anti-nucleocapsid (grey). Values for Relative Intensity Units (RIU) are shown on the y-axis. Cut-off values for a positive test result are shown by the dotted line with the corresponding color. **B**. Three different formulations were used for immunization. In Group 1, two animals received NTD-Fc antigen. In Group 2, two animals received RBD-Fc antigen. In group 3, two animals received a mixture of the NTD-Fc and RBD-Fc antigens. The number of doses received are shown by syringe icons next to the animal ID. **C**. The timeline shows the scheduling of the prime immunization, boosts, and biweekly blood draws. The third immunization at 6 weeks was only administered to animal P0001 as shown. Made in Biorender.com.

The purified proteins were formulated with TiterMax® Gold, an adjuvant selected for its ability to elicit a strong antibody response, but low toxicity and injection site trauma in animals^33^. The immunization of the macaques using the NTD-Fc and RBD-Fc recombinant proteins was expected to produce an antibody response targeted to those regions of the Spike protein, but the number of doses required to elicit a response was unknown. Therefore, all animals received a primary immunization at the beginning of the study and a boost at 2 weeks with a third dose planned for 4 weeks (Fig. 2B, C). Serum from the immunized animals was collected 14 days after the first immunization, prior to the second immunization. This 2-week sample was screened for the presence of antibodies using the CSA: SARS-CoV-2 assay. The standard procedure for this assay uses serum diluted 1:100. The relative levels of anti-spike IgG were evaluated against an established cut-off value for a positive test result.

Spike-specific antibodies were detected in all animals (Fig. 3A). Indeed, the levels of anti-spike IgG in serum from 2 weeks post-immunization reached the upper limits of quantification for the assay (Fig. 3A). The antibody response for group 1, which received the NTD-Fc antigen, generated a detectable response at week 2 but not as strong as groups 2 and 3 (Fig. 3A). Interestingly, both groups 2 and 3 reached the maximum level of detection for the serum concentration tested (1:100 serum dilution) after a single immunization (Fig. 3A). These results demonstrate that a robust IgG response was generated using both the NTD-Fc and RBD-Fc immunogens and that a single dose may be sufficient to generate a robust antibody response.

**Figure 3.**
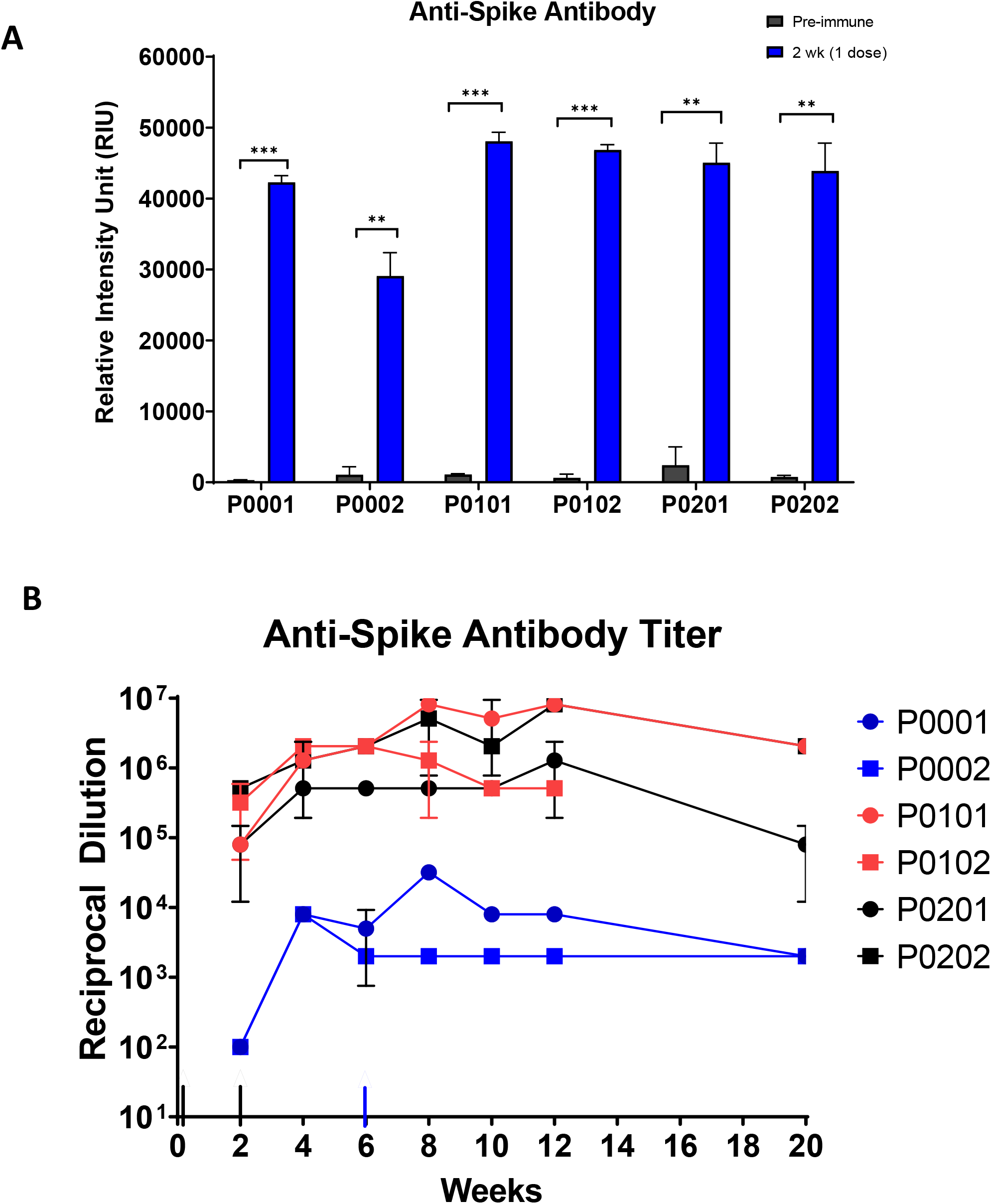
Immunization Antibody Profiles. **A**. Comparison of anti-spike S1 IgG before and after primary immunization. Diluted serum (1:100) from pre-immunization (0 weeks) and 2 weeks post-immunization are compared for antibodies against spike S1. ** p-value ≥0.01, *** p-value ≥0.001. **B**. Antibody titer over 20 weeks. Sera from 2 to 20 weeks post-immunization were titered using the CSA: SARS-CoV-2 assay. The titer was determined as the dilution at which the sample has a signal above 3 standard deviations from the mean of the pre-immune sera. Black arrows indicate the timing of the prime and first boost immunization; blue arrow indicates the second boost administered only to P0001. Error bars show the standard deviation of duplicate measurements.

Following the second immunization, 5 of the 6 animals exhibited local injection site reactions upon routine evaluation, and as a result, further immunizations were halted. Only animal P0001 in group 1 received a third immunization at week 6, based on the comparatively low antibody response at weeks 2 (Fig. 3A) and a lack of local injection site reaction.

### Endpoint spike binding antibody titers in immunized macaques

To better understand the levels and duration of the antibody response to vaccination, endpoint antibody titers were determined for each serum sample collected throughout the study (Fig. 3B). Clear differences in antibody response were observed between group 1 that received the NTD-Fc alone and groups 2 and 3 that received either the RBD-Fc alone or the RBD-Fc/NTD-Fc antigen mixture. Indeed, sera from groups 2 and 3 all had IgG titers greater than 1×10^5^ through 20 weeks post-immunization while sera from group 1 had antibody titers of approximately 1×10^4^ or less throughout the study. Animal P0001, that received a third NTD-Fc dose at week 6, had a higher antibody titer at 12 weeks post-immunization compared to animal P0002 that received only two doses of the same antigen. Nevertheless, both NTD-Fc immunized macaques had antibody levels that were orders of magnitude lower than the other treatment groups. These data suggest that immunogens containing the RBD perform better than the NTD-containing antigens at eliciting a strong and lasting IgG response.

### Inhibition of RBD-ACE2 binding by immune sera

To further investigate the antibody response to the candidate vaccines, we examined the ability of sera from immunized macaques to inhibit RBD binding to ACE2. This binding assay (Intuitive Biosciences, Madison, WI) utilizes ACE2 immobilized in microarray format that is then incubated with recombinant RBD-Fc in solution. If an interfering molecule is present, such as antibodies that prevent binding of the spike RBD to ACE2, a decrease in signal is observed. The commercially available neutralizing mouse monoclonal antibody 5B7D7 was used as a positive control and exhibited the expected inhibition of ACE2 binding (Fig. 4A).

**Figure 4.**
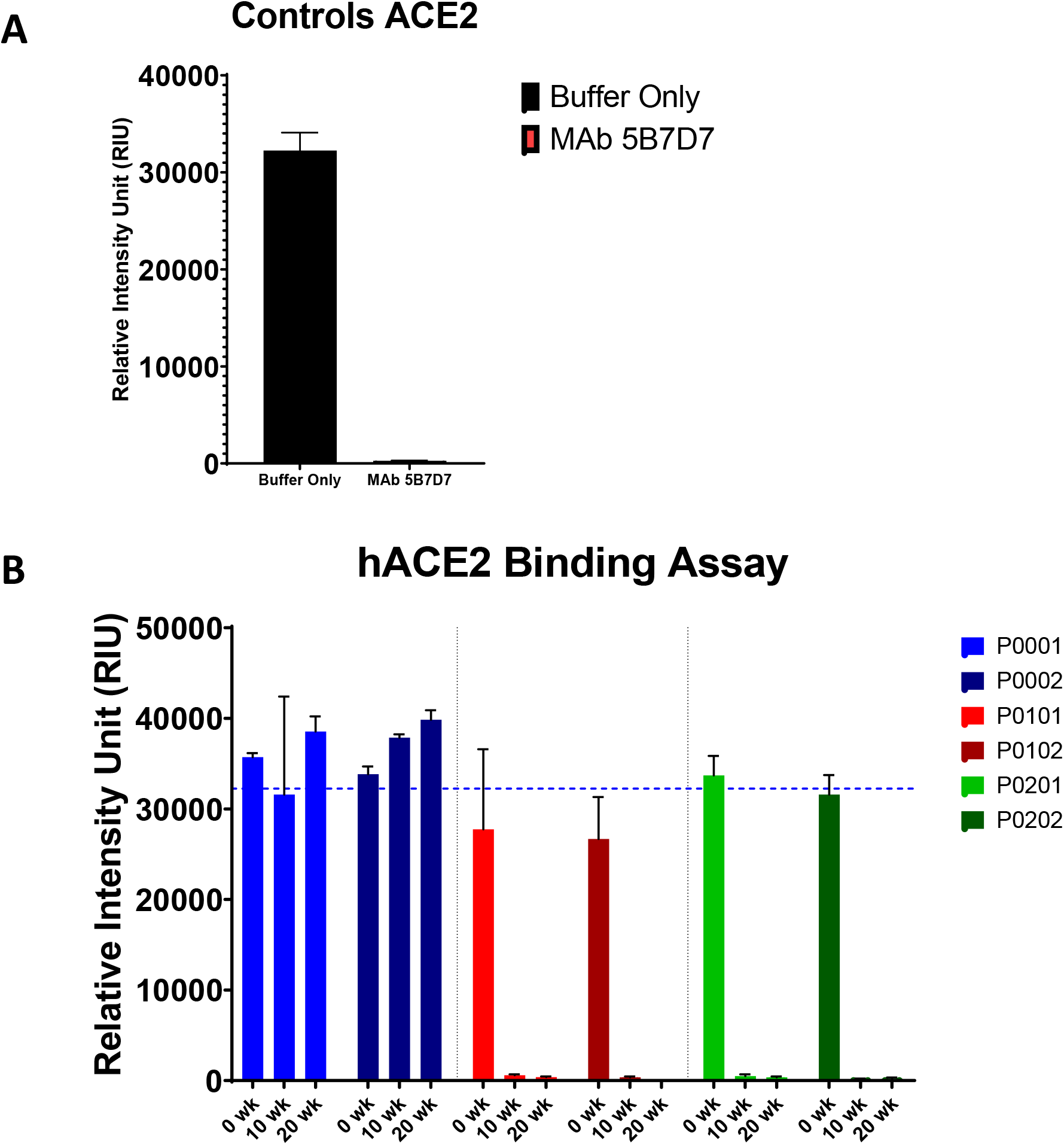
RBD-hACE2 Binding Assay. **A**. Binding of soluble SARS-CoV-2 recombinant spike RBD to immobilized recombinant human ACE2 (hACE2) is measured by Relative Intensity Units (RIU) on the y-axis. Maximum RBD-ACE2 binding is shown in the Buffer Only column (black). Pre-incubation of RBD with the neutralizing monoclonal antibody 5B7D70.1 demonstrates inhibition of hACE2 binding (red). **B**. Sera from 0,10- and 20-weeks post-immunization were assayed for inhibition of RBD binding to hACE2 using a 1:100 dilution. Blue dotted line indicates maximum RBD binding (buffer alone). Bars plotted show the mean RIU in triplicate measurement for animals immunized with NTD-Fc (blue), with RBD-Fc (red), and NTD-Fc + RBD-Fc (green). Error bars indicate the standard deviation of triplicate measurements.

As demonstrated in Figure 4B, sera from group 2 and 3 animals inhibited binding of RBD-Fc to immobilized ACE2. In contrast, sera from group 1 animals did not inhibit RBD-ACE2 binding in this assay. Indeed, because only recombinant RBD was used in this binding assay, antibodies generated from NTD-immunized animals were not expected to interfere with binding to ACE2. Overall, these data show that immunization with RBD-Fc elicits antibodies in nonhuman primates that inhibit RBD binding to ACE2.

### Neutralization of SARS-CoV-2 S-pseudotyped virus by immunized sera

We next determined whether immunization with the NTD or RBD constructs would produce neutralizing antibodies. Neutralization assays were performed using a replication-defective single-cycle pseudotyped virus carrying SARS-CoV-2 spikes and the NanoLuc luciferase reporter. This assay has been previously shown to accurately predict serum neutralizing activity against authentic SARS-CoV-2^24^. As a control for neutralization sensitivity, we used human serum obtained from a SARS-CoV-2 negative individual alone or spiked with a monoclonal neutralizing antibody (Fig. 5A). Serum samples collected at the various timepoints from 0 to 20 weeks post-immunization were evaluated for neutralizing activity. Sera from animals in groups 2 (P0101, P0102) and 3 (P0201, P0202) had readily detectable neutralization activity, as early as 2 weeks post-immunization that significantly increased until weeks 4 to 8 of the study (Fig. 5A, B). Indeed, neutralizing titers were exceptionally high at 4-8 weeks after immunization, in the range of 10,000 to 100,000 and were maintained in the 1000 to 10,000 range at 20 weeks after immunization. In contrast, serum samples from animals in group 1 had undetectable neutralization activity at all time points (Fig. 5 A, B). Comparison of the neutralization titers obtained from group 2 and 3 samples revealed that inclusion of the NTD-Fc did not result in increased neutralizing antibody levels compared to the RBD-Fc alone (Fig. 5 A, B). Additionally, halving the amount of RBD-Fc used did not appear to affect the levels of neutralizing antibodies that were elicited.

**Figure 5.**
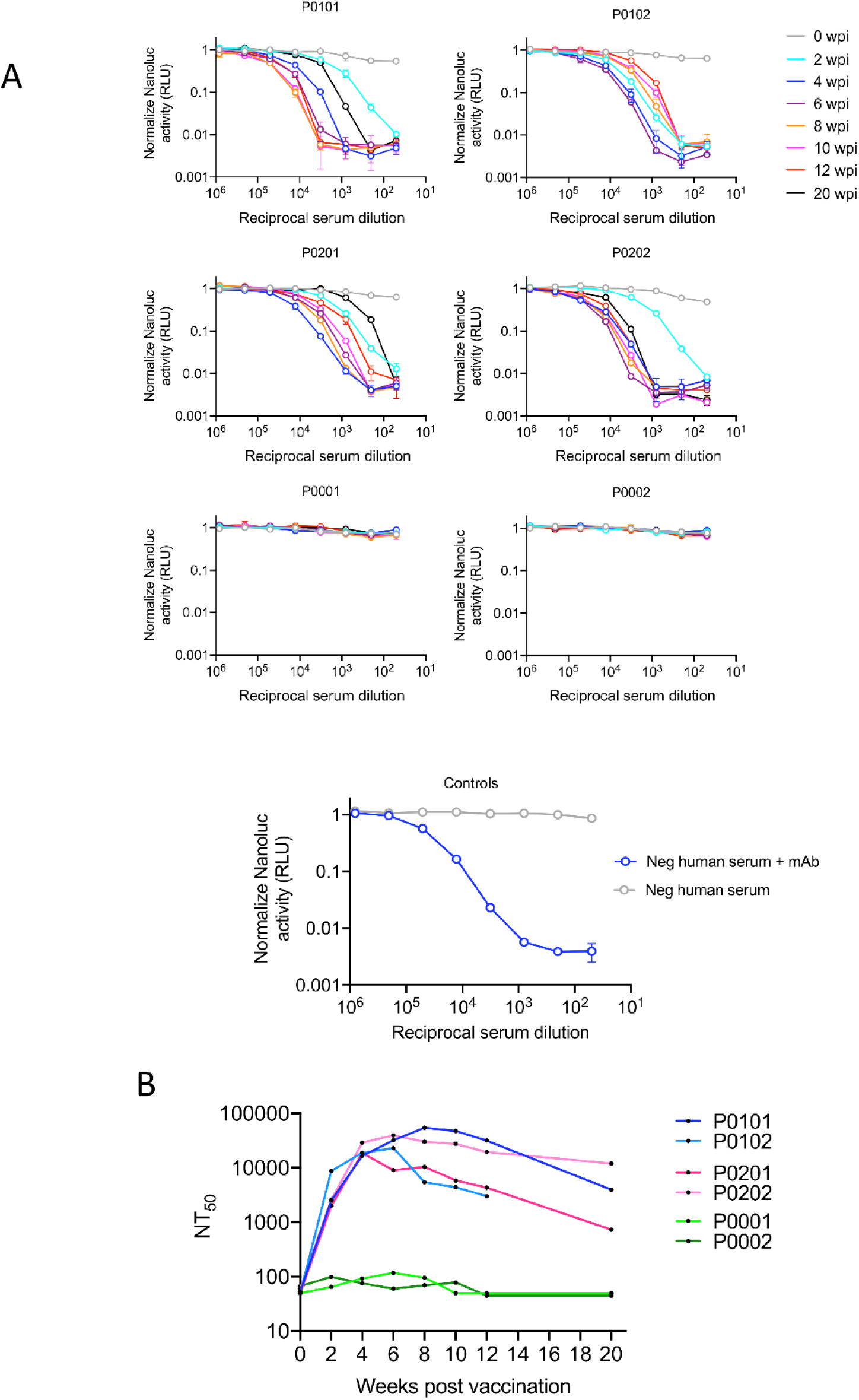
Neutralization of HIV-1 SARS-CoV-2 pseudotype viral infectivity by immunized serum. **A**. Neutralization assays using serum samples from Group 2 (P0101, P0102), Group 3 (P0201, P0202) or Group 1(P0001, P0002)). Infectivity of serum neutralized pseudotyped virus using 295T/ACE2 target cells was quantified by measuring NanoLuc luciferase activity (RLU) and graphed on the y-axis. Reciprocal serum dilution is shown on the x-axis. **B**. The 50% neutralization titer (NT50) for plasma samples shown (A) in a plotted against time post immunization.

## Discussion

The development of highly immunogenic and safe vaccines will be critical for controlling the COVID-19 pandemic, and no one type of vaccine will likely fill the global need. Here, we describe the selection of immunogens that are effective at generating high titers of neutralizing antibodies for sustained periods of time and their evaluation in a NHP model species.

While the COVID-19 pandemic is caused by SARS-CoV-2, prior research on related coronaviruses have revealed key features that shaped vaccine and therapeutic development efforts. A key immunological target is the viral surface protein that is required for virus entry and replication ^34^. Indeed, the spike protein has been the primary target for SARS-CoV vaccines and has been demonstrated to elicit a robust and protective immune response in nonhuman primate animal model systems^29^. Thus SARS-CoV-2 vaccine development has benefited from experience gained in the SARS-CoV vaccine studies. These studies demonstrated that neutralizing antibodies inhibit the interaction of spike protein with its receptor ACE2 and therefore prevent viral entry and replication^17; 32; 35^.

Neutralizing antibodies from recovered COVID-19 patients target epitopes of spike S1, more specifically the RBD and NTD^18; 36; 37 15; 17; 38^. Studies by Qi *et al*. have shown that an RBD-Fc fusion protein (residues 319-541 fused to a mouse IgG1 Fc domain) elicited potent neutralizing antibodies in mice^39^. A conflicting report from Wang *et al*. showed in a murine model that immunization with the RBD expressed with a norovirus shell domain was not sufficient to elicit neutralizing antibodies and the entire S1 portion was needed^40^. While the RBD-based constructs are similar to the one used in this study, the RBD-Fc fusion protein used here also contains an additional 50 residues (541-591) of the C-terminal S1 domain, which may aid in immunogenicity directly, by providing more potential epitopes, or indirectly, by enhancing stability of the RBD-Fc construct or by providing conformational flexibility. Our C-terminally extended RBD construct dimerizes through the disulfide bridge present in the Fc portion, which may also provide non-linear epitopes in its dimerized form. Most proteins benefit from the enhanced pharmacological stability and solubility provided by fusion to a Fc domain^41^. In addition, an Fc fusion can increase uptake by antigen presenting cells (APC) that express the Fc receptor, which in turn can enhance immunogenicity^42; 43^. Our findings in the macaque model system are consistent with the work of Qi *et al*., as we found that immunization with the extended RBD-Fc containing formulations resulted in the generation of high antibody titers (Fig. 3B) with the ability to inhibit RBD binding to ACE2 (Fig.4) and robust neutralizing activity (Fig 5B).

A prior study by Ren *et al*., which used whole spike S1-Fc as an immunogen in macaques and a complex immunization schedule with five doses and multiple adjuvants, reported the successful generation of neutralizing antibody titers^44^. We sought to test an immunization schedule with fewer administrations. The study was originally designed to deliver a primary immunization followed by 2 boosts, 2 weeks apart if needed. However, it was determined that the first 2 doses elicited a sufficiently robust antibody response with endpoint titers exceeding 10^6^ (Fig. 3C) in the RBD-Fc receiving group (2 and 3) and strong neutralizing activity (Fig. 4C, 5A).

While studies have shown SARS-CoV-2 infection of humans elicits antibodies that target the NTD^16; 18^, immunization with the NTD-Fc elicited only a moderate antibody response with a maximum spike-binding titer that was several orders of magnitude less than the extended RBD-Fc. Moreover, no neutralization activity was detected at any time (Fig. 5A). While this does not rule out a possible role for the NTD as an immunogen, it indicates that our extended RBD-Fc construct is sufficient for elicitation of a strong neutralizing antibody response. Using the mix of the RBD-Fc and NTD-Fc (group 3) resulted in the generation of a similar level of neutralization activity as did the use of the RBD-Fc alone (group 2) (Fig. 5B), indicating that the NTD-Fc does not contribute to the neutralizing activity in this context. These data also suggest that a lower dose of the extended RBD-Fc may be as effective as a higher dose, because group 3 received 50% less RBD-Fc than group 2. In addition, the neutralizing antibody titers generated by the extended RBD-Fc immunogen was significantly higher than is typical of individuals who have recovered from a natural SARS-CoV-2 infection or have been immunized with an mRNA or adenovirus vaccine^19; 45; 46; 47^. In preclinical studies in rhesus macaques immunized with the mRNA-1273 vaccine, a reduction in both spike binding IgG and neutralizing antibodies was observed at 8 weeks after the initial vaccination^48^. This suggests that the RBD-containing C-terminal domain protein used in this study (RBD-Fc) may result in a stronger, more durable antibody response than the newly approved mRNA-1273 vaccine.

Clearly an immunogen that can elicit a robust and durable antibody response would be a preferred vaccine candidate. In this study, we demonstrated a robust and sustained humoral response to immunization but did not examine the cell mediated response, which may contribute to protection. Studies with recovered COVID-19 patients have identified additional factors beyond spike-mediated binding and entry that may be important for protection or clearance of virus^49; 50; 51^. However, studies by Du *et al*. on SARS-CoV demonstrate that protection from viral challenge was largely due to neutralizing antibodies resulting from RBD immunization^52; 53; 54^. Further studies using the extended RBD-Fc and NTD-Fc could examine additional aspects of immunity, alternative adjuvants, alternative dosing schedules including a single dose in macaques.

## Conclusions

Overall, this work demonstrates that the SARS-CoV-2 spike extended RBD-Fc fusion protein is effective at eliciting a potent neutralizing antibody response in a macaque model. This response remains high past 20 weeks after the first immunization and is consistent with prior work demonstrating the high antigenicity and robust humoral response elicited by RBD-Fc vaccination. Further studies with the extended RBD-Fc protein would seek to further refine this subunit-based vaccine candidate for optimization of delivery, safety, and efficacy.

## Materials and Methods

### Recombinant Proteins and Antibodies

For vaccination, two recombinant proteins representing portions of the spike S1 subunit were generated. Using Genbank entry MN908947 (identical to NC_045512.2), the predicted amino acid sequence for the spike S1 (aa1-638) was generated using nucleotides 21,563 to 23,476. CHO-cell codon optimization and DNA synthesis was performed at Twist Bioscience (South San Francisco, CA). High-fidelity PCR amplification of coding sequences using Q5 DNA polymerase (NEB, Ipswich, MA) was performed for regions encoding the N-terminal domain (NTD) (Spike protein aa16-309) and the receptor bind domain (RBD)-containing C-terminal domain (CTD) (Spike protein aa319-591) of S1. Coding sequences were cloned using standard molecular biology methods into a Lytic Solutions LLC proprietary expression plasmid to generate secreted S1 fragments fused to a C-terminal human IgG1 Fc. Resulting plasmids were verified by DNA sequencing. Fc-fusion proteins were expressed via transient transfection of suspension-cultured CHO-S (Invitrogen/ThermoFisher) cells using the Mirus Bio, LLC (Madison, WI) CHOgro® High Yield Expression System. Recombinant protein was affinity purified from cell culture medium using Protein-A agarose (Lytic Solutions). Protein was buffer-exchanged into HEPES-buffered saline, quantified, and analyzed by SDS-PAGE for purity. The resulting recombinant proteins are designated as NTD-Fc (16-309) and RBD-Fc (319-591).

The Colony Surveillance Assay CSA: SARS-CoV-2 kit was used (Intuitive Biosciences, Madison, WI) to measure antibodies against recombinant Spike S1 (aa16-685), Spike S2 (aa686-1213), and Nucleocapsid (aa1-419) by multiplex immunoassay on the CSA platform. This product uses SARS-CoV-2 recombinant proteins expressed in CHO or HEK cells to ensure proper glycosylation.

For the human ACE2 (hACE2) binding assay, an RBD-Fc fragment (319-541) with a C-terminal human Fc-tag was purchased from BPS Bioscience (San Diego, CA). Recombinant biotinylated ACE2 was purchased from Acro Biosystems (Newark, DE).

### Animals

Six adult male cynomolgus macaques (*M. fascicularis*) of Asian mainland origin were group housed in European guideline (ETS 123) compliant pens (2 animals/pen) at Covance Laboratories (Greenfield, IN) an AAALAC-accredited facility. At the initiation of dosing, all animals had been given a washout period from previous treatments of at least 56 days. Animals ranged from 48 to 75 months of age and weighed 5.6 to 9.6 kg and were determined to be in good health condition following general clinical observations, blood chemistry and hematology, food consumption, and body weight evaluations.

All animal procedures were approved by Covance-Greenfield Institutional Animal Care and Use Committee and were in compliance with the Novartis Animal Welfare Policy. All procedures adhered to and were in compliance with the Animal Welfare Act and the Guide for the Care and Use of Laboratory Animals, and in compliance with the Office of Laboratory Animal Welfare’s Public Health Service Policy on Humane Care and Use of Laboratory Animals.

### Vaccination

The animals were randomized into three groups (n=2) and scheduled for immunization and serial blood draws over the 20-week study.

NTD-Fc or RBD-Fc proteins were diluted in sterile PBS and formulated as an emulsion with TiterMax Gold adjuvant (Sigma-Aldrich, St. Louis, MO) to a final antigen concentration of 250 µg/mL. For the group dosed with the NTD-Fc/RBD-Fc mixture, each protein was included at a 125 µg /ml concentration to make a final antigen concentration of 250 µg/ml. Formulations were prepared fresh on the day of dosing and used within 3 hours of preparation.

Animals were dosed intramuscularly at 0.8 mL (200 µg), divided into 0.4 mL injections in the quadriceps muscle, per animal. All groups were dosed on Days 1 and 14. For animal P0001 a third NTD-Fc dose was administered on Day 43. For all six animals, blood draws were scheduled at the beginning of the study and every 2 weeks for 12 weeks. For 5 of the 6 animals, one additional blood draw was performed at 20 weeks. Blood was collected from a femoral vein into serum separator tubes (SST). The collected SST were centrifuged within 1 hour of collection for 10 minutes in a refrigerated centrifuge at 2700 rpm. Serum was aliquoted and stored at -80°C until use.

### Serology Testing

Serum samples were evaluated using the multiplex antigen Colony Surveillance Assay (CSA): SARS-CoV-2 kits from Intuitive Biosciences for detection of antigen specific antibodies. SARS-CoV-2 assay plates were manufactured by dispensing picoliter droplets of recombinant antigens and assay controls (BSA-gold, human IgG, buffer only) in a grid-like pattern on the bottom of CSA protein binding plates. Blocking was performed with CSA Buffer and plates dried prior to use. To perform the assay, samples were diluted 1:100 in CSA Buffer and incubated in the antigen microarray well for 1 hour at room temperature. Following 3 washes, a 1:200 dilution of anti-simian gold conjugate was incubated for 1 hour at room temperature. Following 3 washes, signal was developed by a 3-minute incubation with SilverQuant(tm) reagents at room temperature. Gold-catalyzed silver deposition signal develops sites where the anti-simian IgG gold conjugate is bound to sample IgG interacting with antigens on the surface. After rinsing each well with water, individual wells were analyzed on the AQ 1000 analysis system (Intuitive Biosciences). The image capture system used a CCD camera to visualize silver deposition and quantify the signal on a white to black scale measured as Relative Intensity Units (RIU). Samples with an RIU above the product cut-off value were considered antibody positive. Cut-off values were established by the manufacturer validation of the assay using 1:100 serum dilution.

To determine endpoint titers serum samples were diluted in CSA Buffer provided with the CSA: SARS-CoV-2 kits. A fourfold serial dilution was performed starting with 1:500 in CSA Buffer to a maximum dilution of 1:8×10^7^. Each dilution was assayed using the CSA: SARS-CoV-2 kit as described above. The highest dilution at which the sample was 3 standard deviations above the mean of the pre-immune value at a 1:500 dilution was considered the antibody titer.

### Spike-hACE2 Binding-inhibition Assay

Neutravidin at 100 µg/mL (Thermo Fisher Scientific, Waltham, MA), BSA-gold (Intuitive Biosciences, Madison, WI), and human IgG (Jackson Immunoresearch, West Grove, PA) were printed with picoliter droplets and immobilized on the surface of microarray plates (Intuitive Biosciences). Biotinylated recombinant human hACE2 (SinoBiological, Wayne, PA) was bound to the neutravidin spots by incubating a 0.1 µg/ml solution of hACE2 in CSA buffer for 1 hr. The plates were wash 3 times with plate wash buffer to remove unbound hACE2 and used immediately. Test serum (1:100 dilution) or the mAb 5B7D7 (1 µg/ml) (GenScript, Piscataway, NJ) was diluted in CSA buffer and incubated for 1 hour at room temperature with 0.1 µg/mL RBD-Fc (BPS Bioscience, San Diego, CA). Pre-incubated samples were individually added to wells of the hACE2 multiplex plates and incubated for 1 hour at room temperature. After washing, a 1:200 dilution of anti-IgG gold conjugate (Intuitive Biosciences) was added to each well and incubated for 1 hour at room temperature to detect the Fc fragment of the RBD-Fc. After washing, the signal was developed with a 3-minute incubation with SilverQuant reagents. The wells were rinsed, dried, and analyzed on the AQ 1000 analysis system (Intuitive Biosciences, Madison, WI). Results were calculated as a percent inhibition of three no serum (CSA buffer only) control wells to establish a maximal binding of RBD-Fc to hACE2 in this assay system.

### SARS-CoV-2 pseudotyped reporter virus and pseudotyped virus neutralization assay

SARS-CoV-2 pseudotyped particles were generated as previously described^24^. Briefly, 293T cells were transfected with pHIV-1NLGagPol, pCCNG/nLuc and pSARS-CoV-2-SΔ19. Particles were harvested 48 hours after transfection, filtered and stored at -80°C. Fourfold serially diluted serum from immunized macaques was incubated with SARS-CoV-2 pseudotyped virus for 1 h at 37 °C. The mixture was subsequently incubated with 293T/ACE2cl.22 cells (plated on Poly-D-Lysine-coated 96-well plates) with the final starting dilution of serum being 1:50. At 48 h later the cells were washed with PBS and lysed with Luciferase Cell Culture Lysis 5× reagent (Promega). Nanoluc Luciferase activity in lysates was measured using the Nano-Glo Luciferase Assay System (Promega) with the Modulus II Microplate Reader (Turner BioSystems). The raw nanoluc luciferase activity values (relative luminescence units) were normalized to those derived from cells infected with SARS-CoV-2 pseudotyped virus in the absence of serum or a rabbit monoclonal antibody diluted in normal human serum at 0.105 mg/mL (40592-R001, Sinobiological, Wayne, PA). The half-maximal inhibitory concentration for serum (NT50) was determined using four-parameter nonlinear regression (GraphPad Prism).

### Statistics

All statistical analyses were carried out with GraphPad Prism software. P < 0.05 was considered significant. Multiple unpaired two-tailed t-test was used to compare the antibody reactivity in pre-immune compared to 2-week post-immunization for each animal, using baseline subtracted values for the buffer only control.

## Declarations

### Ethics approval and consent to participate

Not applicable.

### Consent for publication

Not applicable.

### Availability of data and materials

The datasets used and/or analyzed during the current study are available from the corresponding author on reasonable request.

### Competing interests

KL and is an employee and shareholder of Intuitive Biosciences. GB is an employee of Intuitive Biosciences. FS is a founder and employee of Lytic Solutions. DR is an employee of Lytic Solutions. DC and JF are current or former employees of Covance Greenfield Laboratories. KM, MB, and LM are employees of Novartis. The remaining authors declare no conflict of interests.

### Funding

Work at the Rockefeller University was supported by grants from NIAID R01 AI078788 (to TH) and R37AI64003 to (PB).

### Authors’ contributions

GB, DR, FM, JF, and DC performed experiments and analyses. FM provided performed and analyzed the neutralization assays. JF, DC, LM, MB, KM provided animal resources, care, and expert counsel. KL supervised the study and wrote the manuscript. TH and PB supervised the neutralization work, revised, and edited the manuscript. This manuscript has been read and approved by all authors.

## Acknowledgements

We would like to acknowledge the contributions of Lindsey Moser, who contributed to the development of the CSA: Simian SARS-CoV-2 assay, during her time working with Intuitive Biosciences.

